# How does selection for flowering in maize shapes defenses components against the European Corn Borer ?

**DOI:** 10.64898/2025.12.12.694016

**Authors:** Sacha Revillon, Inoussa Sanane, Matthieu Reymond, Valérie Méchin, Nathalie Galic, François Rebaudo, Fréderic Marion-Poll, Christine Dillmann, Judith Legrand

## Abstract

Understanding how selection for flowering time affects maize susceptibility to the European corn borer (ECB, Ostrinia nubilalis) is essential in the context of climate change and increasing pest pressures. We investigated the consequences of divergent selection for flowering time on maize defense and susceptibility traits using two genetic backgrounds (MBS847 and F252). Three complementary approaches were combined: leaf palatability tests, ECB field incidence dynamics surveys, and biochemical analyses of maize stem walls. In the MBS847 background, early-flowering families exhibited lower content of defense compounds, higher leaf palatability, and higher field infestation compared to late-flowering families. In contrast, no significant differences were observed between early and late populations in the F252 background. Differences in susceptibility between families were associated with variation in stem biochemical composition, including fiber components which influence stem rigidity and resistance to ECB. Our results indicate that selection for flowering time can modify plant defense traits and susceptibility in a genetic-background–dependent manner. These findings highlight the interaction between plant phenology, biochemical defenses, and insect herbivory, with implications for breeding and adaptation under changing environmental conditions.

## 1 Introduction

Domesticated plants are generally more susceptible to herbivory than their wild relatives due to the selection of traits during domestication that favor production, such as number of fruits or size of fruits, and palatability such as high nutritional content (Whitehead et al., 2017; Moreira et al., 2018; Solís-Montero et al., 2020; Fernandez et al., 2021). Several mechanisms could explain such susceptibility including direct selection on traits related to plant defense (e.g. toxicity, taste, nutritional content) or indirect selection impacting plant defenses such as pleiotropy or allocation costs (Whitehead et al., 2017; Fernandez et al., 2021; Chen et al., 2015). To maximize their fitness, plants must perform three essential functions: growth, reproduction, and defense (Stamp, 2003; Cipollini et al., 2017; Züst and Agrawal, 2017; Watts et al., 2023). In resource-constrained environments, investing in one of these functions may limit resources allocated to the others. For example, setting up defenses involves synthesizing compounds or organs, which requires resources that could not be allocated to growth (He et al., 2022). Due to such trade-off, breeding for higher growth and yield would result in a drop of defense levels. In maize, the plant model used in this study, teosinte has been shown to have better defense mechanisms than modern maize, but it also has a lower yield (Rosenthal and Dirzo, 1997; Fontes-Puebla and Bernal, 2020). Domestication has also indirectly impacted maize defense strategies against certain pest species, notably the expression of resistance, i.e. plant’s ability to negatively impact herbivores, and tolerance, i.e. plant’s ability to reduce herbivore impact on fitness. It has been shown that maize resistance to Western corn rootworm decreased with domestication and its spread whereas tolerance increased (Whitehead et al., 2017; Fernandez et al., 2021; Fontes-Puebla and Bernal, 2020). An increase in tolerance could result from breeding for high yield, as plants must be able to limit yield losses due to herbivory. The associated decrease in resistance may occur due to a trade-off between tolerance and resistance (Salgado-Luarte et al., 2023). This is in line with the plant defense hypotheses suggesting that plants cannot maximize both defense strategies simultaneously due to limited resources (Stamp, 2003). During the 20th century, breeding programs were implemented to reintroduce resistance in maize by developing inbred lines exhibiting either high chemical or physical defenses (Russell et al., 1975; Willmot et al., 2004). Moreover, some selection programs have been carried out on criteria indirectly linked to defenses, such as vegetative development traits, to exploit the potential consequences of resources allocations trade-offs (Revilla et al., 2005; Riedeman et al., 2008; Sandoya et al., 2010; Ordas et al., 2013). Some of these studies target traits that are not correlated with yield in order to maximize defense and productivity (Revilla et al., 2005; Riedeman et al., 2008). For example, Revilla et al. (2005) investigated the effect of the timing of transition from the juvenile to the adult vegetative phase in maize for three cycles of selection on the damages of European and Mediterranean corn borers (ECB and MCB) under artificial infestation. Results showed no significant effect of the selection on the length of tunnels made by borers into the stem and ear damages. Similarly, Riedeman et al. (2008) conducted a comparable selection on seven generations and evidenced no differences on leaves damages and the number of infested plants. Other studies have investigated the effect of selection for flowering time on plant susceptibility to ECB and MCB as earliness was shown to be positively correlated with resistance (Samayoa et al., 2014), although negatively correlated with yield (Ordás et al., 2019). Understanding these relationships is crucial in the context of global warming with emerging questions about crop ability to adapt to future growing conditions (Olesen et al., 2012; Challinor et al., 2016; Dong et al., 2021), while facing increasing pest outbreaks (Bebber et al., 2013). Adjusting crop flowering time could therefore represent a key strategy to avoid high-risk periods due to climatic hazards and intense pest pressure (Slot et al., 2021). In areas affected by climate change stress, farmers have already begun adapting their cultural practices by sowing crops earlier or by using early-maturing varieties to reduce crop damage caused by climate uncertainty (Shaffril et al., 2018). It is therefore crucial to better understand the effect of selection on flowering time on plant susceptibility to pest in order evaluate the viability of such adaptation of these methods for crop safety. Ordas et al. (2013) studied the resistance to ECB and MCB (tunnel length, ear damages) of several cycle of selection for early flowering and showed that early flowering genotypes suffered less from artificial infestation than late flowering genotypes. Tissue maturity at the onset of artificial infestation was the limiting factor for pest infestation, with early flowering genotypes being the most developed. Overall, the relationship between flowering time and susceptibility is not yet fully understood. In this study, we examined the effect of selection for flowering time on plant susceptibility by investigating the relationship between plant earliness, plant defense components, and pest damages in the field. The original plant material consisted of genetically related maize inbred lines derived from two divergent selection experiments (DSE) for flowering time conducted for over 20 years on the Plateau de Saclay, France (Durand et al., 2010). For each DSE, starting from a single inbred-line seed lot, one early-flowering and one late-flowering inbred line populations have been produced by successive self-fertilization of early or late flowering plants (Durand et al., 2010). Selection for flowering date has led to significant changes in plant phenology, with marked shifts in the date of floral transition (Tenaillon et al., 2018). The specific selection regime applied to Saclay’s DSE, with a combination of small population size and a strong selection intensity, was shown to specifically enrich the selected lines with genes involved in the genetic determination of maize phenology (Desbiez-Piat et al., 2021). Therefore, when comparing early and late lines derived from the same ancestor, the main difference is expected to be the plant phenology or traits that evolved along the flowering time selection process. Using this plant material, we made use of the annual implementation of the Saclay’s DSE experimental design to evaluate several defense traits against ECB that naturally infests maize fields in the Plateau de Saclay. To determine whether differences in ECB susceptibility are related to plant earliness, we conducted three experiments comparing parental lines with early and late populations derived from selection. First, we grew a subset of late and early-flowering lines from the 18th generation to measure their palatability to ECB larvae (Sanane et al., 2021). Second, we sampled plants from the 23rd generation and quantified a set of traits related to cell-wall digestibility using both biochemical assays and near-infra-red spectrometry (El Hage et al., 2018). Third, we monitored the dynamics of natural field infestation during two successive years, 2018 and 2019, within the DSE experimental set-up, corresponding to the 22nd and 23rd generations of the DSE. Combining these approaches, we tested the effect of selection for flowering time on various maize susceptibility traits.

## 2 Materials and methods

### 2.1 Materials

#### 2.1.1 Plant material

The plant material is issued from the Saclay Divergent Selection Experiments (Durand et al., 2010). These experiments were initiated from two inbred lines, F252 and MBS847, in order to produce, within each genetic background, two populations differing for their flowering time (hereinafter referred to as early and late populations). F252 is an American flint, and MBS847 is a late iodent dent obtained in 1992 from the breeding companies Agri-Obtentions (F252) and Mike Brayton Seeds (MBS847). In each genetic background, each of the two populations (early and late) was initiated from two distinct plants of generation zero, thereby giving rise to two independent families per population (Figure S1). At each generation, in the early families, the ten earliest flowering plants were selected and self-fertilized, while in the late families, the ten latest flowering plants were selected and self-fertilized. A selected plant is hereafter referred to as a progenitor. For the F252 genetic background, the early families are named FE036 and FE039, and the late ones FL317 and FL318 (F stands for F252, E for early, and L for late). For the MBS847 genetic background, the early families are named ME049 and ME052, and the late ones ML040 and ML053 (M stands for MBS847, E for early, and L for late), overall eight different families were evolved from the two initial seed lots. Plants are thus structured into genetic backgrounds (MBS847 or F252), populations within genetic backgrounds (early or late), families within populations (two families per population), and progenitors within families. In the following, we refer to each family derived from the divergent selection experiment, regardless of its genetic background (F252 or MBS847) or its population (early or late), as an inbred line. Inbred lines from three generations (G18, G22, G23) of the DSE were used in this study (Figure S1). Four families from generation G18 (FE036, FL317, ME052, and ML053) were sown in the field for the palatability experiment in 2018 and 2019. For the ECB field infestation dynamics survey, we used the plants sown for the annual DSE experiment from generation G22 (2018) and G23 (2019). Biochemical analyses were carried out on plants sampled from the eight families of the generation G23 in 2019.

#### 2.1.2 Insect material and rearing

The artificial population of corn borers (*Ostrinia nubilalis*) used for the palatability tests was provided by the Bioline AgroSciences breeding farm (France). Eggs were reared at 26±1°C until larval emergence, with a (16:8) (day: night) photoperiod, at 70% humidity. After emergence, larvae were fed with an artificial diet (composition 1.32 L water, 27 g agar, 224 g corn flour, 60 g dried yeast, 56 g wheat germ, 12 g ascorbic acid, 4 g vitamin mix (Ref. 0155200), 0.8 g Aureomycin/tetracycline, 2 g Nipagin, 1.6 g sorbic acid and 4 g benzoic acid). Larvae aged from 10 days after emergence were used for palatability tests. For field incidence monitoring, *O. nubilalis* populations were those naturally infesting the experimental site.

### 2.2 Experimental settings

#### 2.2.1 Experimental design of the DSE

For both years, the field in which DSE were carried out was divided into twenty-two (in 2018) and nineteen (in 2019) blocks of sixteen plots. In each plot, twenty-five plant seeds were sown. The first and last blocks of the field were borders as well as the first and last plots of each block. All the inbred lines derived from the same ancestral line (either MBS847 or F252) were sown in eight contiguous blocks. Within each background, blocks sown with early inbred lines alternated with blocks sown with late inbred lines. In each family, plants were issued from four, five or six different progenitors (Figure S1). For each progenitor, four plots were sown constituting four replicates, with one replicate per block. For each plot, during the flowering period, the number of flowered plants was recorded daily. These daily data were used to compute the median flowering time at the plot level in thermal time. Thermal time was computed with the thermal time function described in Parent et al. (2010) using the daily mean temperature data collected from the field meteorological station (Delannoy et al., 2022). Additionally, the plant height of all self-fertilized plants was measured as the distance from the ground to the bottom of the tassel. These data were used to compute an average height per plot.

#### 2.2.2 Measuring inbred lines biochemical traits

In order to study the cell wall composition of the inbred lines of generation G23 of the DSE in 2019, biochemical analyses were carried out on non-infested plants within the field of DSE, at the silage harvest stage. For each progenitor, two of the four sown plots were randomly selected, and three plants per plot were sampled. For each plant, the green aerial plant parts were collected from the base of the aerial roots to the tassel and ears were removed. Each sample was ground and dried in an oven at 60°C for 48 h (Zhang et al., 2019). The dry matter powder was analysed at the Institut Jean-Pierre Bourgoin according to the protocol used by (El Hage et al., 2018). Biochemical analyses and Near-Infra Red Spectroscopy (NIRS) were carried out to measure or estimate thirty-two variables, including cell wall constituents and whole-plant digestibility (Table S1).

#### 2.2.3 Measuring inbred lines palatability for ECB larvae

In 2018 and 2019, plants from the G18 generation of the DSE were sown in a separate field trial to assess their palatability for reared ECB second instar larvae (L2) at different dates. The palatability test is described in detail in (Sanane et al., 2021). Briefly, it consists in monitoring leaf discs palatability by ECB larvae (2nd instar), and inferring palatability behaviours, from the most to the least consumer. One replicate consisted in a 50-well (5×10) plate with 50 leaf discs (one disc per well) sampled from the last ligulated leaf from two to three plants of the same inbred line sampled at the same date. Each plate was first filled with agarose 1% to prevent the leaf discs from drying out. Plates were transported from the field to the lab, and palatability tests were realised immediately after the field sampling. A single L2 instar larva was placed on each leaf disc and its palatability was monitored for 48 hours, using a webcam generating image stacks. Image stacks were analysed using the RoitoRoiArray and Areatrack plugins of the Icy imaging software (De Chaumont et al., 2012). The plugins were used to delineate pits and leaf discs in a semi-automated approach and then to track changes over time in the proportion of leaf disc consumed (number of pixels in each leaf disc). Image analysis generates measurements (pixels per minute) for each leaf disc in each well, which are exported as text-based files for analysis. The data were then converted into TSV flat files.

The palatability of four families, one early and one late inbred line derived from each parent used in the DSE (ME052, ML040, FE036, FL318) was assessed at four different dates in 2018 and six dates in 2019 (Figure S2). The sampling dates were the same for all inbred lines. In 2018, plants from the same plot were used for all sampling dates. In 2019, plants from different plots were used at the different sampling dates. Two replicates per inbred line and per sampling date were used, except for FL317 and ME052 that had only one replicate for all sampling dates in 2018 and for the first sampling date in 2019.

#### 2.2.4 Measuring ECB field infestation dynamics of inbred lines

We monitored the natural field-infestation dynamics of ECB on the DSE field for two consecutive years (generation G22 in 2018 and generation G23 in 2019). To avoid destructive observations, a plant was considered as infested when at least one ECB-typical damage was observed on the aerial parts of the plant. The types of damages varied along the plant and the insect development: typical holes on leaves, stems, tassels and ears, typical broken stems or tassels.

Weekly observations were performed from July 5 to September 20 in 2018 and from July 12 to September 24 in 2019. In 2018, for each plot, the number of plants showing damages was counted weekly. In 2019, observations were done at the plant level. Each week, newly attacked plants were marked using coloured bands corresponding to the week of observations. Once marked, plants were not observed anymore in the following weeks. From these observed data, the number of attacked plants per plot on a given week was computed to obtain a measure comparable to the 2018 observations. For each year, each plot and each week, the incidence was computed as the ratio between the number of plants showing damages and the total number of plants on the plot.

### 2.3 Statistical analysis

#### 2.3.1 Studying the effect of selection on cell-wall composition

In order to study the effect of selection on plant defense traits, biochemical analyses of stem walls of inbred lines of the generation G23 were carried out in 2019. Biochemical analysis data were grouped with plant growth traits data (plant height and phenology) into a single dataset for statistical analysis. All measured variables were taken at the plot level represented by a pool of three plants for biochemical traits, the median of the plot for flowering time and the mean of the three self-fertilized plants for the height. The variations of each studied trait were modeled using a linear mixed effect model:

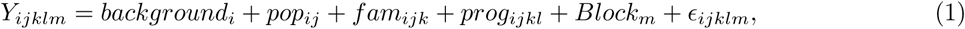

where *Y_ijklm_* corresponds to the trait value for the plot of the progenitor *l*, from the family *k*, in the population *j* (early or late), in the genetic background *i* (MBS847 or F252) in the block *m*. The background, population (pop), family (fam) and progenitor effects were modeled as nested fixed effects. The Block effect was modeled as a Gaussian random effect of variance 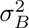, and *ɛ_ijklm_* were Gaussian residuals of variance *σ*^2^. Maximum likelihood tests were used to compare nested models, and to test the significance of genetic background, population, family, and progenitor effects. The test of the Block variance component was performed using the varTestnlme package (Baey and Kuhn, 2023). Trait means of each progenitor estimated with model (1) were used to identify which biochemical traits differentiate early and late families within each genetic background with a Factorial Discriminant Analysis (FDA).

#### 2.3.2 Identifying variability in inbred lines leaf palatability

To study the relationship between leaf palatability and earliness, leaf palatability tests were carried out on four progenitors representing four families from the generation G18 of the DSE in 2018 and 2019. The statistical methods that were used to analyse the data have been described in (Sanane et al., 2021, 2023). Briefly, a replicate consisted in a plate of 50 leaf discs producing 50 consumption curves. Using the SOTA algorithm (Herrero et al., 2001), all curves were classified into 14 typical profiles. Then, k-means were used to classify these profiles into six consumption behaviour groups from the most (A) to the least consumer (F) (Figure S3). Consumption behaviours were then reduced into two classes by grouping the most consumer groups into one single consumer group (C) and the least consumer groups into one single non-consumer group (N). To quantify the palatability for one replicate, the *CNratio* was defined as:

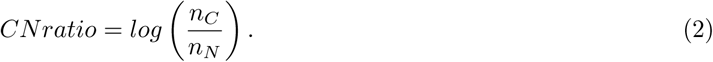

with *n_C_*the number of C curves in the replicate and *n_N_*the number of N curves in the same replicate (*n_C_*+ *n_N_* = 50).

The composition of groups C and N from groups A to F was chosen in order to maximise the coefficient of discrimination of a model explaining the CNratio by the year, the family and the plant development state.

In order to consider growth rate variability between inbred lines, we quantified plant development state of a plot at each monitoring date as the thermal time difference between the sampling date and the average flowering date of the plot. A negative value indicates a sampling date before flowering. A positive value indicates a sampling date after flowering. Notice that as the two replicates from the same maize line were taken in the same plot at a given sampling date, there is a single average development state for the family by sampling date combination. The average development state observed for each sampling date and each maize line is represented in Figure S2. The following statistical model was used to study the effect of the plant development state on the *CNratio*:

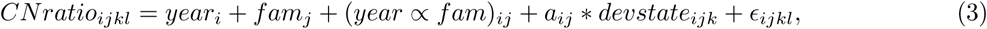

with *CNratio_ijkl_*the *CNratio* of family *j*, on year *i*, on date *k* for replicate *l*, *dev_ijk_* the development state, *year_i_*and *fam_j_* respectively the effects of the year and family factors, and *a_ij_* the slope of the linear relation between the *CNratio* and *dev_ijk_* for year *i* and family *j*.

All possible combinations to define groups C and N (*e.g.*, C = *A* and N = *B* + *C* + *D* + *E* + *F*, C = *A* + *B* and N = *C* + *D* + *E* + *F*,…) were tested in model (3). The combination maximizing the R^2^ was C=A+B and N =D+E+F leading to the *CNratio* definition:

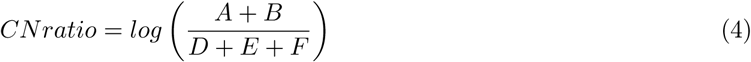

Once a definition of *CNratio* was chosen (Table S4), the different effects in the model (3) were tested comparing nested models.

#### 2.3.3 Identifying variability within inbred lines in field infestation dynamics

In order to quantify the infestation dynamics of ECB in the field, the incidence was computed for each plot throughout the season as the number of infested plants in a plot divided by the number of grown plants in these plots. The variations of incidence according to the inbred line were studied using two statistical modeling approaches: one spatial and one spatio-temporal model. The spatial model, fit separately on the data from each week, is described below, whereas the spatio-temporal model, fit at once on the whole dataset, is described in Supplementary Information, as this approach fails to converge when estimating a progenitor effect. Analyses were conducted on each year and genetic background separately. Spatial auto-correlation due to ECB adult movements and ECB larval dispersion was integrated in the model using a Spatial Temporal Conditional Auto-Regressive model (STCAR) in a Bayesian framework (Lee et al., 2018). Different neighborhood structures were compared as described below.

The incidence of plot *k*, *Y_k_*was modeled as a Gaussian variable with the mean *µ_k_* having two components, the family or progenitor effect *α_k_*and the spatial effect *ϕ_k_*:

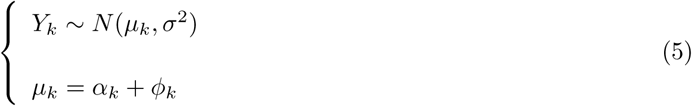

An independent Gaussian prior is specified for *α_k_*. The spatial correlation induced by *ϕ_k_* is defined in Leroux et al., 2000 by (6) with a Uniform prior on *ρ* and *τ* ^2^:

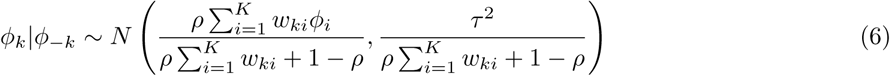

The spatial correlation is weighted by a neighborhood matrix W = (*w_kj_*). W is binary where *w_kj_* = 1 if plots *k* and *j* are considered as neighbors. Three spatial structures between blocks and between plots were tested. The first structure sets *w_kj_* = 1 if plot *k* and *j* are adjacent within a block and 0 otherwise. The second sets *w_kj_* = 1 for all the plots *j* in the block of plot *k*. The third sets *w_kj_* = 1 if plot *j* and *k* are adjacent within a block or if the plot *j* of the surrounding block is adjacent with plot *k*.

Models were fitted using Markov chain Monte Carlo simulations with the R packages CARBayes (Lee, 2013) and CARBayesST (Lee et al., 2018). Regression and variance parameters were sampled using a Gibbs sampler. Random effects and the remaining parameters were sampled using the Metropolis algorithm. For the spatial model fit separately on data from each week, three independent chains were run for 120 000 iterations with a burn-in period of 20 000 and a thinning of 10. First, a visual inspection of the three traces of the Markov chains was made in order to identify some correlations in chains or non-convergence scenarios.

This inspection highlighted a lack of power for the first two weeks of the season. Moreover, the Geweke coefficients of the different Markov chains were used to assess convergence of the different simulations. The deviance information criterion (DIC) (Spiegelhalter et al., 2002) was used to select models with or without spatial auto-correlation, models with different spatial structures, and models with family or progenitor effect. The distribution of the incidence of each genetic background (resp. population or family) was computed by aggregating the Markov chains of incidence of all the progenitors constituting the genetic background (resp. population or family). In this Bayesian framework, we considered that the effect of a factor was significant when the 95% Credible Interval (CI) for one of its parameters did not include zero.

## 3 Results

Three generations (G18, G22, and G23) of maize inbred lines issued from a divergent selection for flowering time in two different genetic backgrounds were grown in agricultural conditions in the Saclay plateau in 2018 and 2019. We quantified different components of plant defenses or pest susceptibility: the biochemical cell wall composition (G23), the palatability of leaf discs for ECB larvae (G18), and the ECB field natural infestation dynamics (G22, G23). First, we describe the plant response to selection for flowering date. Then, we present the effects of selection on each studied trait measured in the three different experimental settings. Finally, we explore how these different components of pest susceptibility relate.

### 3.1 Response to selection on flowering time

Response to selection on flowering time of Saclay DSE was studied for the twenty first generations of selection in (Durand et al. 2010, Durand et al. 2015, Desbiez-Piat et al. 2021), highlighting a strong response to selection. In this work, we used the G18, G22 and G23 generations of selection. For each generation, significant differences were found between flowering dates of the two genetic backgrounds (F252 or MBS847, p-value *<* 0.001 for G18, G22, G23), of the two populations within each genetic background (early versus late, p-value *<* 0.001 for G18, G22, G23), of the two families within populations (p-value *<* 0.001 for G18, G22, G23) and of the different progenitors within families for G22 (p-value *<* 0.001). The estimated means of flowering date in thermal time for each family are shown in Table S2. For both generations G22 and G23, there is approximately a 10 thermal days (corresponding to one day of development at 20°C, according to the temperature response function of Parent et al. (2010)) difference in flowering time between the early and late families of the F252 genetic background, and a 20 thermal days difference between the early and late families of MBS847 background.

### 3.2 Selection for lateness tends to strengthen plant composition and defense

In 2019, plants from the generation G23 of the DSE were characterized by a set of phenotypic measurements including flowering date, plant height, and variables measuring the chemical composition of maize stems at late development states, before the completion of grain maturation (see Table S1).

For each trait, the sources of trait variations were quantified and tested using an ANOVA of model (1). Complete results of the ANOVA are presented in Table S3. Among the 29 biochemical traits, 11 showed significant differences between genetic backgrounds, 21 showed differences between early and late populations within the same genetic background, 14 traits showed differences between families and 3 traits showed differences between progenitors. More specifically, significant differences in plant composition between early and late populations can be observed in compounds relating to lignin (kl_cwr, abl_cwr), its composition (subunits H and G) and stem rigidity (cellulose, hemicellulose), all these compounds being in higher content in late populations within each genetic background than in early populations. Together, selection for differences in flowering time resulted in differences between early and late populations concerning cell wall composition at plant maturity.

A factorial discriminant analysis (FDA) was performed using the means per progenitor estimated with model (1) and setting the progenitor family as the discriminant factor to identify which biochemical traits differentiate early and late families within each genetic background and how they are related. The two first FDA axes explain 95.7% of the total inertia (Figure 1). Within each genetic background, selection for lateness is related to a shift to the right of axis 1. In the MBS847 background, it is also related to a shift to the bottom of the second axis. Within each background, early families are grouped with ancestral lines. The variables most positively correlated with axis 1 are biochemical variables such as klason lignin (kl_ms, kl_cwr), neutral detergent fiber content (KLADL_NDF), percentage of S-subunit lignin content (pS), cell wall residue (cwr), cellulose content (cellulose_ms) and silica content (cendre_ms). In addition, flowering time (flo), presented as a supplementary variable, is highly positively correlated with the first axis. The variables most negatively correlated with axis 1 are H-subunit lignin content (pH) and plant digestibility (IVDMD, IVCWRDm) (Figure 1b). Thus, axis 1 separates families according to their earliness, digestibility and stem rigidity, but also for biochemical variables related to defense compounds (kl_ms, H, pS, abd_cwr). Axis 2 correlates negatively with dry matter content (ms) and acetyl bromide lignin (abl_cwr). Hence, for both genetic backgrounds, selection for lateness is related to an increase in plant rigidity (cellulose_ms, hemicellulose_ms), in klason lignin (kl_ms, kl_cwr) and in silica content (cendre_ms), a decrease in plant digestibility (IVDMD, IVCWRDm), and complex modifications of the composition of lignin subunits (pS, pG, pH). In addition, in the MBS847 background, selection for lateness is related to an increase of the content in some defense compounds.

**Figure 1:**
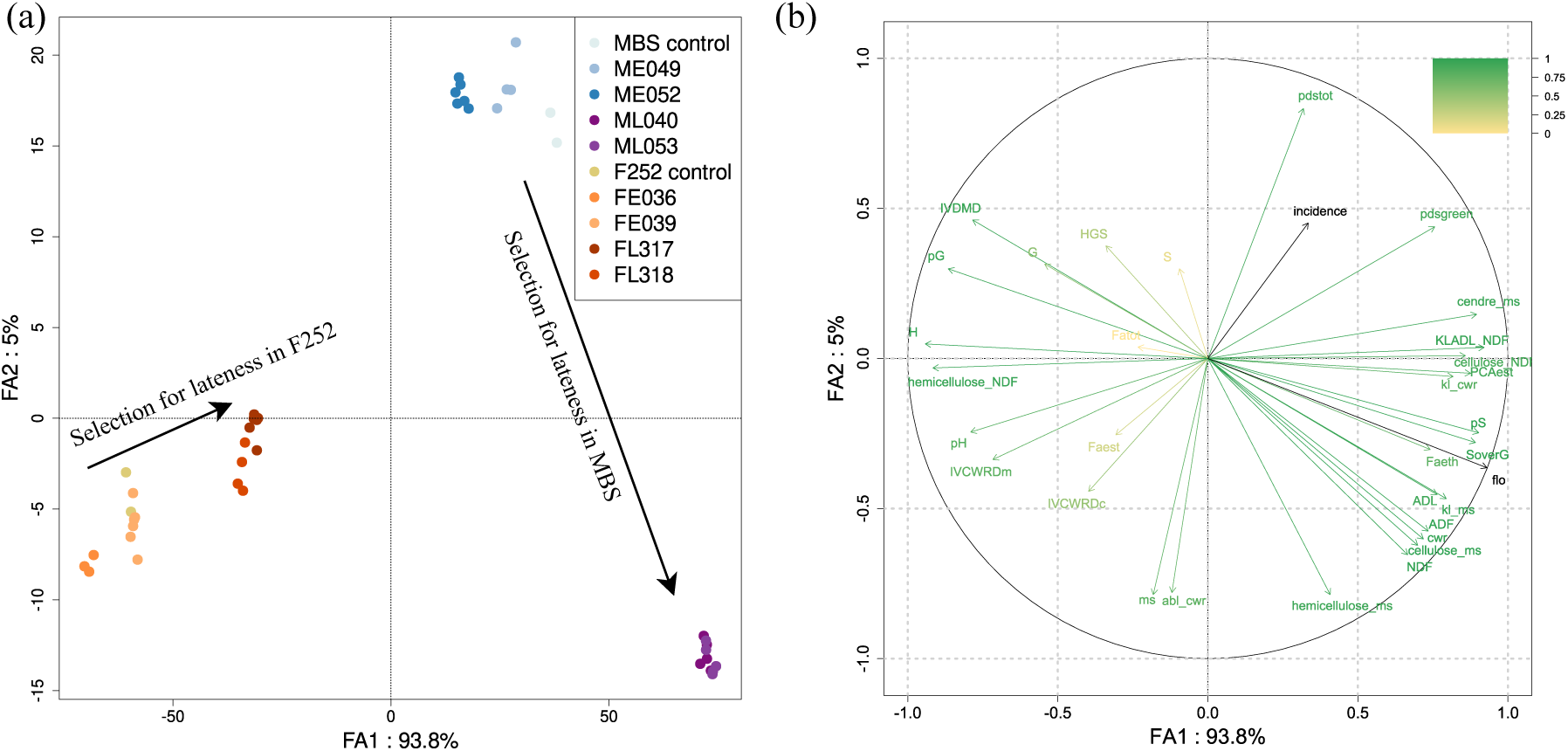
(a) Factorial Discriminant Analysis (FDA). FDA was performed using the estimated means per progenitor by model (1) and setting the plant family as the discriminant factor. Each color represents a family. (b) Correlation circle of variables (listed in Table S1). The variables “flo” (the mean flowering date in thermal time in 2019) and “incidence” (the mean final pest incidence in 2019) are represented as supplementary variables.

### 3.3 Leaf palatability can be affected by selection for flowering time and is mainly determined by plant development state

In order to quantify genetic variations for leaf palatability and investigate its relationship with plant development, ECB leaf consumption behaviours were analysed from plants of generation G18 of the DSE sampled at regular stages throughout their development before and after flowering time. In order to disentangle the effects of sampling plants differing in growth rates, we set up an original measurement of plant development state defined as the difference in thermal time between the sampling date and the time to flowering. As expected, at each sampling date, a development state gradient was observed with early F252 being more advanced than late F252 (on average 2 to 4 thermal days difference), early MBS847 being more advanced than late MBS847 (on average 4 to 8 thermal days difference), and F252 being more advanced than MBS847 (on average 10 to 15 thermal days) (Figure S2). The leaf palatability was quantified by the *CNratio* (high CNratio indicating high palatability, see Methods) and its variations components were analysed with model (3). *CNratio* depends significantly on the plant family (p-value = 4.14e-2) and decreased with development state (p-value < 0.001) with a slope independent from the plant family. For all families, the leaf palatability decreased with plant development (Figure 2). At the same development state, plants from the MBS847 late population were significantly less palatable than plants from the MBS847 early population (p-value = 4.16e-2). In the F252 genetic background, no differences were evidenced between early and late populations (p-value = 0.95) (Figure 2). Plants from the late MBS847 population were less palatable than those from the F252 genetic background (p-value < 0.001), but not less palatable than those from the early MBS847 population. This suggests that selection pressure has modified leaf palatability in the MBS847 genetic background but not in the F252 genetic background.

**Figure 2:**
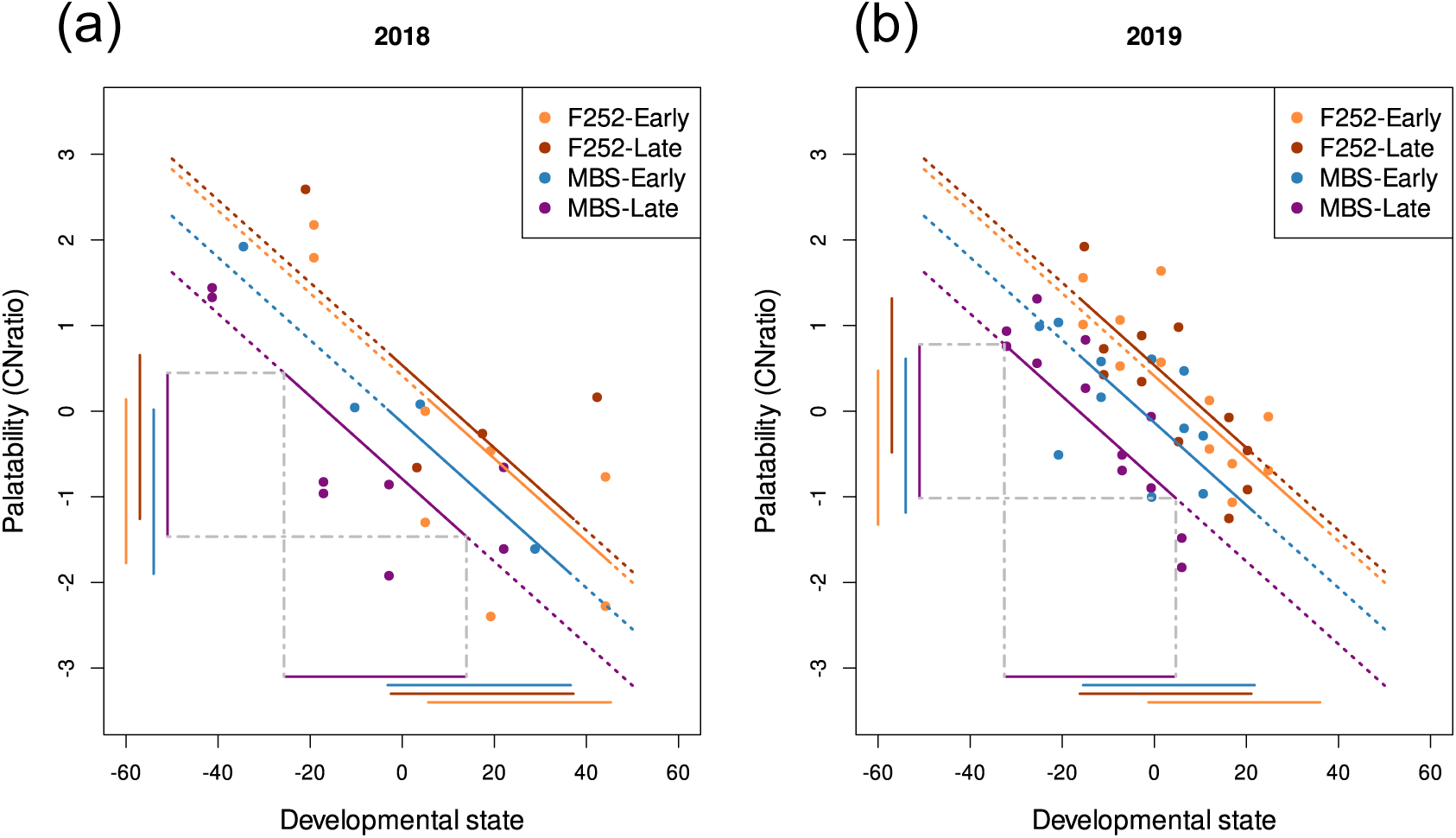
Relationship between CN ratio (a proxy of palatability) and development at each leaf disc sampling date in 2018 (a) and 2019 (b). Colored points represent the observations, and lines represent the model-estimated means (3). The solid portion of each line corresponds to the CN ratio associated with the range of development stages of the plants during the exposure to the first generation of the pest in the field incidence experiment. Horizontal bars indicate the periods of pest pressure, and vertical bars mark the corresponding CN ratio.

### 3.4 ECB larvae favor early MBS847 population in the field

In order to study another component of pest success, ECB infestation dynamics over the season was evaluated by weekly monitoring the infested plants in the field between early July and late September. Each year and each genetic background dataset were analysed separately. For both years and both genetic backgrounds, the model with spatial correlation and the third spatial structure was selected with the DIC criterion. For each week of observation, the spatial correlation *ρ_S_* and its variance *τ* ^2^ were estimated fitting model (5). For both years and each genetic background, the CIs of *ρ_S_* were very broad (Figure S3). The estimated median of *τ* ^2^ was low for all the weeks and did not exceed 1% (Figure S4). Thus, the model (5) did not reveal any clear spatial correlation pattern.

The dynamics of the median pest incidence varied between years and plant families (Figure 3). In 2018, incidence globally increased from the beginning of July to reach a plateau at 30% before suddenly increasing again at the end of August. In mid-September, the median incidence reached almost 100% for some families. In 2019, incidence increased between July 19 and August 23 and reached a plateau at 15 to 30%, depending on the family. A slight increase in incidence was observed at the last monitoring date in September. These observations are consistent with the existence of two successive generations of ECB. A first one in spring, with damages observable on the plants in early July. A second generation at the end of the summer, whose larvae colonized the plants at the end of August in 2018 and mid-September in 2019. Differences in incidence have been evidenced with model (5) between early and late populations within genetic backgrounds (Figure S6 et Figure S7), between families (ME049, ME052, ML040 and ML043 for MBS847 background, and FE036, FE039, FL317 and FL318 for the F252 background) (Figure 3) and between progenitors within families but concerned mainly progenitors of the FE036 family (Figure S8 and Figure S9).

**Figure 3:**
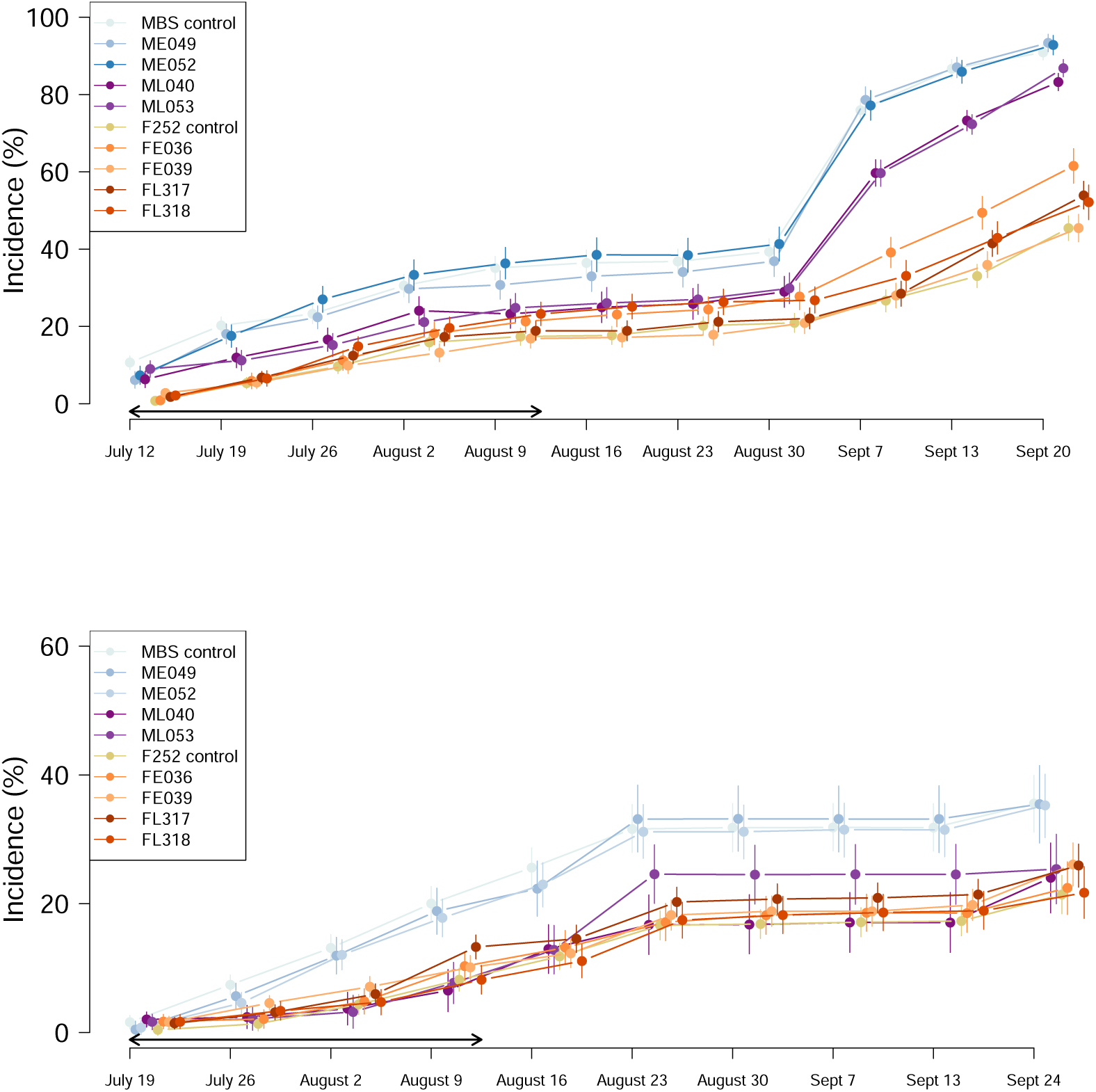
Incidence of borer damages by plant family (2018 above, 2019 below). Abcissa axes are the monitoring date. These results were obtained using the weekly approach and the model with spatial correlation on the MBS847 and F252 data separately. For each family, the average incidence over the progenitors and the three markovs chains is computed. The quantiles at 2.5% and 97.5% and the median are displayed. Each color correspond to a family. Black arrows represent the exposure period reported on Figure 2.

Although model (5) revealed significant differences in incidence between early and late populations, between families and between progenitors in both genetic backgrounds, the patterns observed differed between backgrounds. In the MBS847 background, a clear and repeatable pattern was found: early populations, early families (ME049, ME052) and the ancestral line were systematically more infested than the late populations and late families (ML040, ML053) in both years. In the F252 background, however, no such recurrent pattern was observed across years: for example, the two early families were either among the least (FE036) or the most infested (FE039) in 2018.

### 3.5 Partial consistency between plant composition, leaf palatability and field incidence

The three approaches used in this study target different components of plant susceptibility on different plant generations. ECB field incidence dynamics (G22 and G23) provide a direct measure of pest susceptibility but might depend on the level and timing of the pest pressure in the field while palatability tests (G18) and biochemical analysis (G23) provide measures on different plant defenses components at different developmental states. Below, we examine how consistent are our findings regarding differences between populations quantified by these three approaches in relation with earliness and developmental state differences.

Within the MBS847 genetic background, the three approaches show a coherent pattern. Late families displayed higher levels of rigidity-related compounds (cellulose_ms, hemicellulose_ms, p-value <0.001) and defense-related metabolites (kl_ms, H, abl_cwr, p-value <0.001), and lower digestibility (IVDMD, p-value <0.001) compared to early families and ancestral lines (Figure 1). Some of these biochemical differences correlated positively with field incidence: plants with lower digestibility (IVCWRD: cor = −34%, p-value < 0.05) and defense content (H: cor = −32%, p-value < 0.05, abl_cwr: cor = −43%, p-value < 0.01) were the most infested. Palatability tests on G18 confirmed this trend, with early plants being more consumed than late plants (Figure 2). This consistency suggests that selection for earliness reduced plant defenses in MBS847, leading to both greater palatability and field susceptibility.

In contrast, in the F252 genetic background, we did not evidence differences in biochemical composition, palatability, or field incidence between early and late families. However, when comparing F252 to MBS847 late families, some inconsistencies appear. For a similar development state, F252 plants had a palatability level higher than MBS847 late plants, which could suggest higher field infestation. Yet, field incidence showed that F252 plants were less or equally infested. This apparent contradiction can be explained by considering plant phenology during the main period of infestation (Figure 2). At that time, the “consumer/non-consumer” ratio was equivalent between F252 and MBS847 late plants, suggesting similar effective palatability in the field. Thus, differences in development timing between genetic backgrounds likely modulate the relationship between biochemical defenses, palatability, and field susceptibility.

In summary, the three approaches yield consistent results within MBS847, whereas the apparent inconsistencies observed between MBS847 late and F252 populations could be explained by differences in plant phenology, which may lead to differences in development state at the time when pest exerts pressure.

## 4 Discussion

We studied the effect of selection for flowering time on maize susceptibility to ECB using three different approaches. We conducted leaf palatability tests, ECB field incidence dynamics surveys, and biochemical analyses of maize stem walls in two different genetic backgrounds (MBS847 and F252). Results showed an effect of selection in the MBS847 genetic background for which plants from the early population have lower content of defense compounds, were more consumed, and more infested in the field than plants from the late population while no significant differences were found between the early and late populations within the F252 genetic background.

Our study stands out by simultaneously investigating three different measures of plant defenses, each capturing a different aspect of plant susceptibility to ECB while most works on the effect of selection for growth traits on maize susceptibility to ECB usually focus on damages at harvest (Revilla et al., 2005; Riedeman et al., 2008; Sandoya et al., 2010; Ordas et al., 2013). Damages caused by ECB vary between larval stages and generations due to larval growth and the development of plant organs during ontogeny (Balachowsky, 1966; Campos et al., 2003). Consequently, a susceptibility trait measured at harvest may not fully represent all aspects of plant susceptibility, which can vary across the season, along plant and insect development. In this study, we quantified damages of both the first and the second ECB generations. Incidence in early monitoring dates describes first-generation damages for which larvae mainly feed on leaves while incidence in later monitoring dates describes damages of the later ECB generations feeding on ears and stems (Balachowsky, 1966). Leaf palatability tests mainly describe damages of the first ECB generation but can also describe damages of the first instars of the second ECB generation for leaves sampled collected after flowering. Moreover, biochemical composition of plants at silage stage describes mainly plant defenses against the second ECB generation. Thus, these traits describe several components of infestation and studying their relationship could help to better understand the different components of plant susceptibility. For example, field incidence results from both larval ability to attack the plant after egg hatching and larval migration may depend on leaf palatability and biochemical composition. Within the MBS847 genetic background, the results of the three approaches support a consistent trend as the least defended plants were the most infested in the field and their leaves were the most extensively consumed in the palatability experiment. Within the F252 genetic background, the results of the three approaches also show consistency as no clear pattern was found but a high variability was shown between plant families of this genetic background for the three studied traits.

Both palatability tests, biochemical analyses and field incidence surveys showed an effect of the selection for flowering time on maize susceptibility in the MBS847 genetic background while not in F252. Within the MBS847 genetic background, a clear pattern was observed between plant families issued from the two directions of selection with early families having a lower content of defense compounds, being more consumed, and more infested in the field than late families. Stem biochemical composition is an essential factor in resistance to ECB. Specifically, the content of studied compounds such as fiber (lignin, cellulose, hemicellulose, Neutral Detergent Fiber) and etherified ferulic acid has been associated with stem rigidity, which has been found to be positively correlated with resistance to stem borers (Santiago et al., 2003; Martin et al., 2004; Santiago et al., 2013; Santiago et al., 2016). Our findings indicate that the content of these compounds has significantly increased through selection for flowering time in MBS847 in the direction of lateness, but not in the F252 genetic background. Some research has focused on studying the correlation of these compounds with agronomic traits such as flowering time in breeding programs for resistance (Santiago et al., 2016; Wolf et al., 1993). Concerning fiber concentrations, Wolf et al. (1993) showed that plants selected for high NDF and lignin content flowered later than plants selected for lower NDF and lignin content which is consistent with our results. Concerning ferulic acid, Santiago et al. (2016) performed a divergent selection program for total cell wall diferulate concentration in maize stem and showed that populations selected for high diferulate concentrations flowered earlier and were taller than populations selected for low concentration. In our study, we did not observe any difference in the total concentration of ferulic (Fatot) between early and late populations. However, we observed an opposite effect on the ferulic etherified content (Faeth), whose level increased with selection for lateness in MBS847. Thus, the effect of earliness on the content of compounds related to stem rigidity varies between compounds, highlighting the complexity of the relationship between these two traits in maize. Indeed, flowering is a complex trait influenced by numerous QTLs (Buckler et al., 2009), some of which are co-localized with QTLs associated with resistance traits (Jiménez-Galindo et al., 2019). The selection for lateness in MBS847 may have targeted these co-localized QTLs, but not in F252. Furthermore, the differences in flowering time between early and late populations in F252 are relatively small (approximately 5 days of thermal time), compared to a larger difference of about 15 days of thermal time observed between early and late MBS847 families. These smaller differences in flowering time within F252 may be insufficient to produce a significant modification in cell wall composition.

In addition to biochemical analyses of plant composition, we documented the infestation dynamics of ECB by monitoring field incidence over the season. When comparing the results of our study with findings in the literature, it is essential to consider that our plant material allows discussion of our results on two different scales. On one hand, we can discuss the results obtained in each genetic background and compare the early and late flowering plant families in MBS847 and in F252. Within each genetic background, the plant families differ for earliness but share a common genetic basis, allowing us to set aside potential genetic resistance factors when making comparisons. On the other hand, we can discuss these results by putting the two genetic backgrounds into perspective and question the relationship between earliness and plant susceptibility at a broader scale. Overall, plants from the F252 genetic background flowered earlier than plants from the MBS847 genetic background and we found that they were less infested than MBS847 plants in the first year of experiment and as infested as the MBS847 late population in the second year of experiment, MBS847 early population being the most infested each year. In the literature, studies have examined the relationship between plant susceptibility and earliness both in selection programs for flowering time (Ordas et al., 2013) and within the genetic diversity of maize (Beres and Gorski, 2012). On one hand, Ordas et al. (2013) have shown that early varieties were less infested, which aligns with our results when comparing F252 and MBS847 genetic backgrounds, but is not in line with our results within the MBS847 genetic background. On the other hand, Beres and Gorski (2012) showed that late varieties were less infested, which this time does not align with our results when comparing F252 and MBS847 but is in line with our findings within the MBS847 genetic background. Several hypotheses could explain these contradictions and help to clarify the relationship between plant earliness and susceptibility. First, we can suppose that such differences may be due to genetic resistance factors between plant materials of the different studies. Indeed, in our study we showed that there was a partial decoupling between content in plant defenses and earliness when comparing MBS847 and F252 genetic backgrounds. Moreover, differences in biotic conditions between the different studies, such as pest voltinism and phenology, may explain such contradictions. It has been shown that the impact of earliness on maize susceptibility can vary depending on pest voltinism (Balachowsky, 1966), which may explain the contradictory findings reported in the literature (Ordas et al., 2013; Beres and Gorski, 2012). Balachowsky (1966) highlighted that for univoltine infestations, early varieties could be more susceptible to egg-laying by the European Corn Borer (ECB) as in (Beres and Gorski, 2012) where both pest voltinism and results coincide. In contrast, for bivoltine infestations, late varieties could be more attractive for egg-laying of adults of the ECB second generation, which causes the most significant damages as in (Ordas et al., 2013) where both pest voltinism and results also coincide. This suggests that pest voltinism could impact the relationship between plant earliness and susceptibility, leading to different findings. In a field experiment, Revillon et al. (2024) studied the effect of maize phenology and development on maize susceptibility to the European corn borer. A panel of 23 maize inbred lines differing for phenology were sown at three different planting dates to expose the pest to various stages of maize development. They demonstrated that plant susceptibility depended on the earliness of the inbred line, the sowing date and their interactions, but also that the effect of earliness on pest infestation could vary depending on pest phenology. More specifically, they showed that the timing of infestation onset relative to plant development impacted plant susceptibility, suggesting an effect of the timing between maize and corn borer phenologies. This could partly explain the apparent inconsistency between the stems biochemical composition analysis, the palatability tests, and the field incidence survey regarding the differences between the MBS847 late population and the F252 genetic background. Indeed, F252 plants show similar content of most defense compounds as the MBS847 late population but were more consumed for the same development state in the palatability test. One might expect, therefore, that F252 would have been more infested in the field. However, in the field, F252 ancestral lines were less or equally infested than MBS847 late populations. But, if we take into account the state of development of the plants during the main period of infestation in the field, we can observe that the “consumer–non-consumer” ratio measuring the palatability of each of the genetic backgrounds was equivalent for the plants in the field. This suggests that the F252 plants and MBS847 late population exhibited both similar levels of leaf palatability during infestation and content of defense that could explain the similar levels of infestation in the field.

In this study, we investigated the effect of selection for flowering time on susceptibility to the European corn borer within a divergent selection experiment in two genetic backgrounds. Selection impacted susceptibility in one of the two genetic backgrounds, where early families exhibited lower levels of defense compounds, were more consumed and more infested in the field compared to late families. These results contribute to the literature on the complex relationship between earliness and susceptibility to the European corn borer in maize. In the context of climate change, which alters crop growing periods and necessitates the adaptation of crop phenology, our results provide valuable insights into the potential consequences of crop adaptation on their resistance to pests, whose outbreaks are predicted to increase (Tobin et al., 2008; Minoli et al., 2022; Katsaruware-Chapoto et al., 2017).

## Notes

### Competing Interest Statement

The authors have declared no competing interest.

