## Supplementray Figures and Tables for "How does selection for flowering in maize shapes defenses components against the European Corn Borer ?"

### Supplementary Information

December 12, 2025

### Supplementary Figure S1

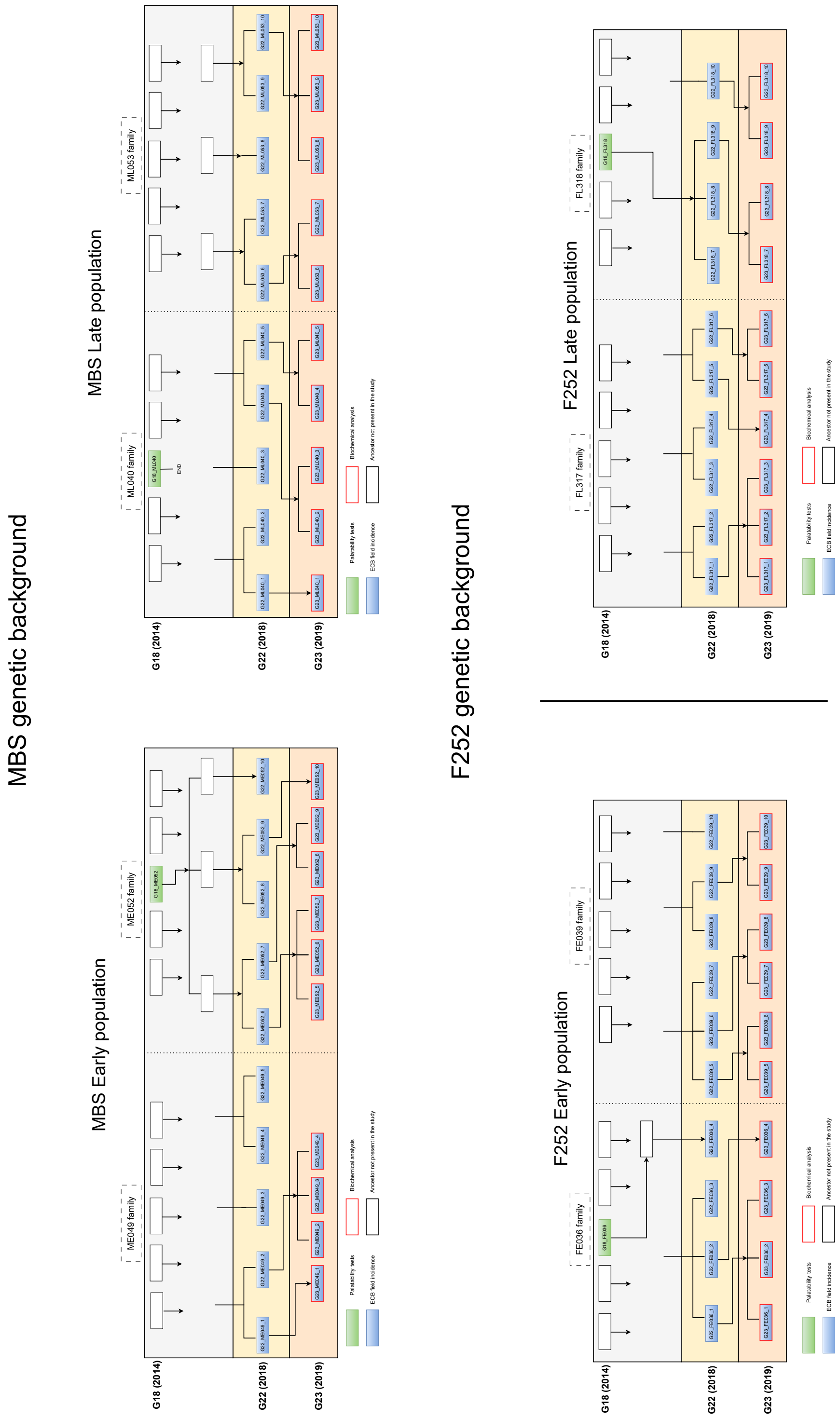

Figure S1 : Plant pedigree of the SDSE. Each box represents a progenitor of either the G18, G22 or G23 generation of the SDSE. Progenitors colored in green were used in the palatability tests. Progenitors in blue were used in the ECB field incidence survey. Progenitors surrounded in red were characterized with biochemical analysis.

#### Supplementary Figure S2

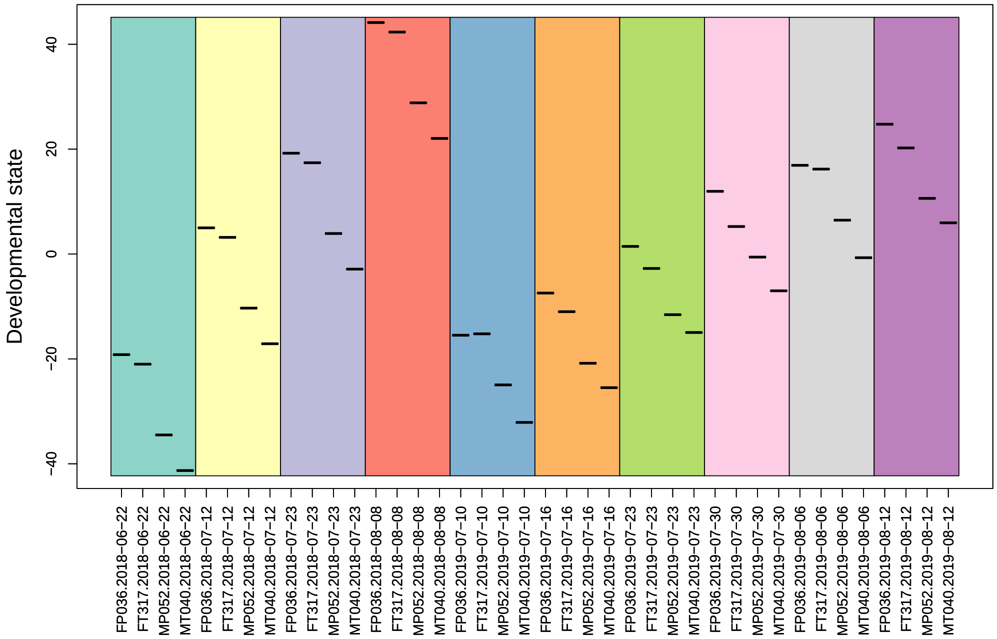

Figure S2 : Plant development stage at each sampling date for the palatability tests. Plant development state was defined as the thermal time difference between the sampling date and the average flowering date of the plot.

### Supplementary Figure S3

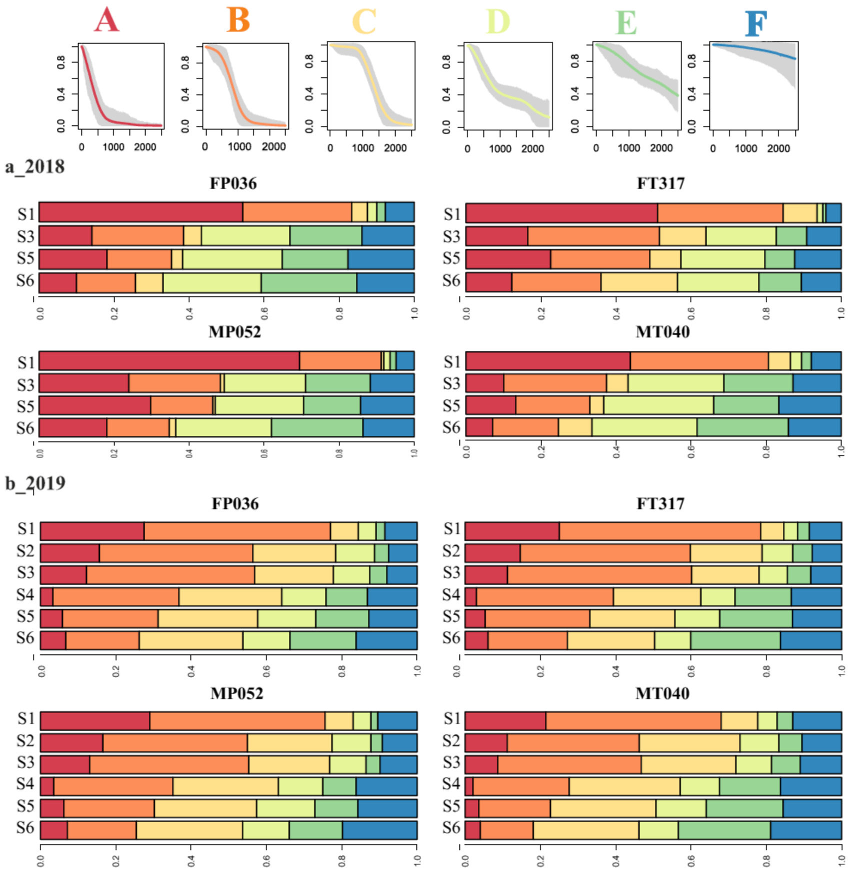

Figure S3. Distribution of feeding behaviors according to the developmental stage of the plants for each family. The estimated proportions of each type of feeding behavior [A–F] are shown in the barplot. The barplots are organized according to the sampling stage from S1 to S6. **a.** Overview of the average consumption dynamics for each type of feeding behavior; the shaded area represents the variability in consumption within each type. **b.** 2018 data. **c.** 2019 data

#### Supplementary Figure S4

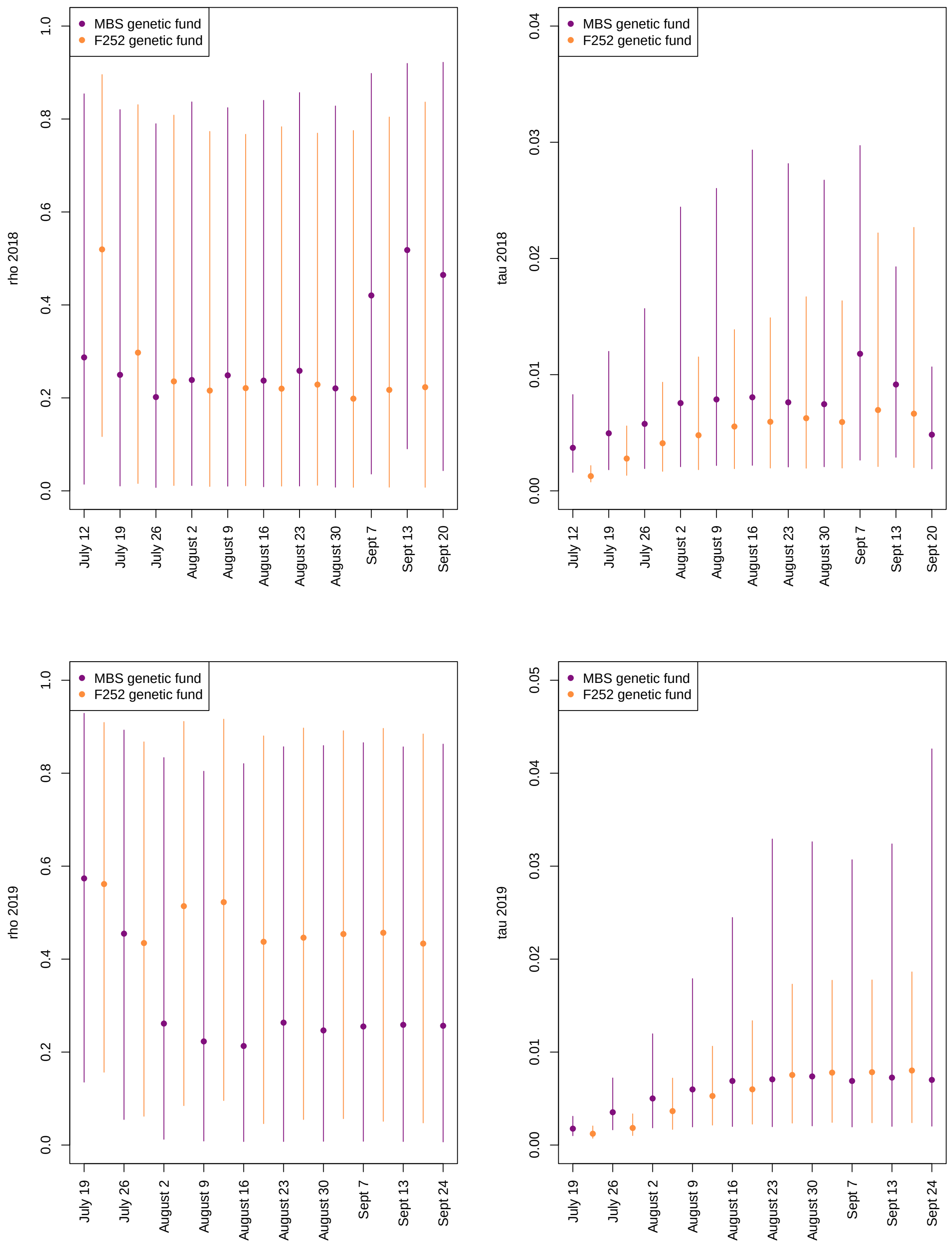

Figure S4 : Estimated median of $\rho_S$ and $\tau^2$ with model (6) within the genetic backgrounds MBS and F252 in 2018 (up) and 2019 (bottom). Error bars represent 95% credible intervals for estimates.

### Supplementary Figure S5

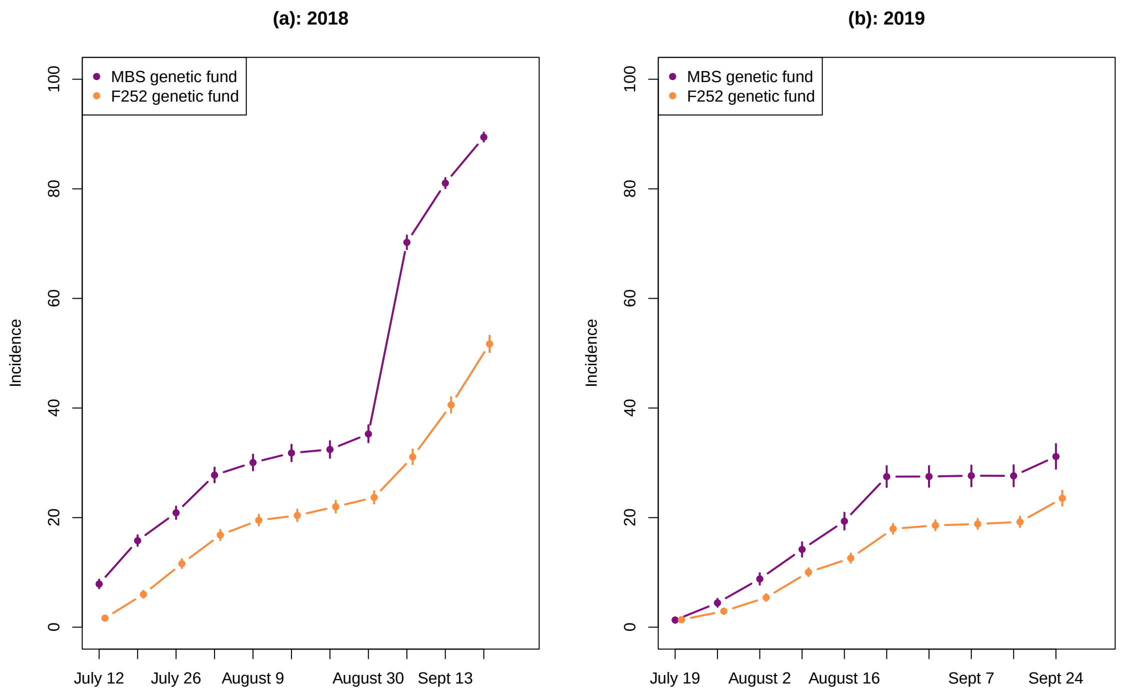

Figure S5 : Estimated medians of the ECB field incidence for each genetic background MBS847 and F252 at each monitoring date in 2018 (a), and 2019 (b). Error bars represent 95% credible intervals for estimates.

#### Supplementary Figure S6

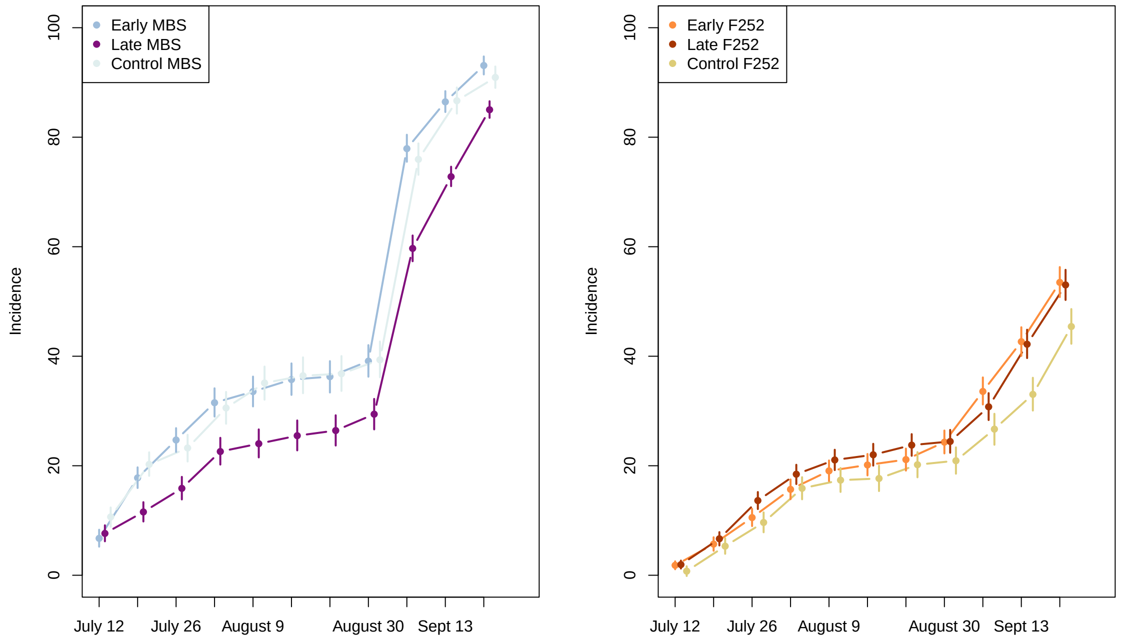

Figure S6 : Estimated medians of the ECB field incidence for each population within the genetic backgrounds MBS847 and F252 at each monitoring date in 2018. Error bars represent 95% credible intervals for estimates.

### Supplementary Figure S7

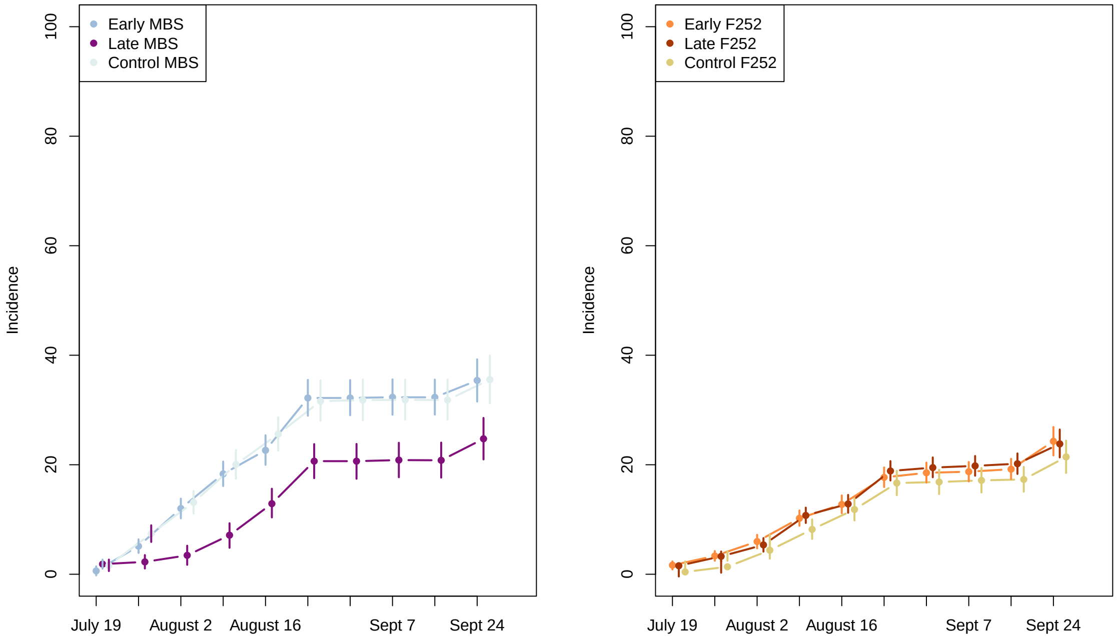

Figure S7 : Estimated medians of the ECB field incidence for each population within the genetic backgrounds MBS847 and F252 at each monitoring date in 2019. Error bars represent 95% credible intervals for estimates.

### Supplementary Figure S8

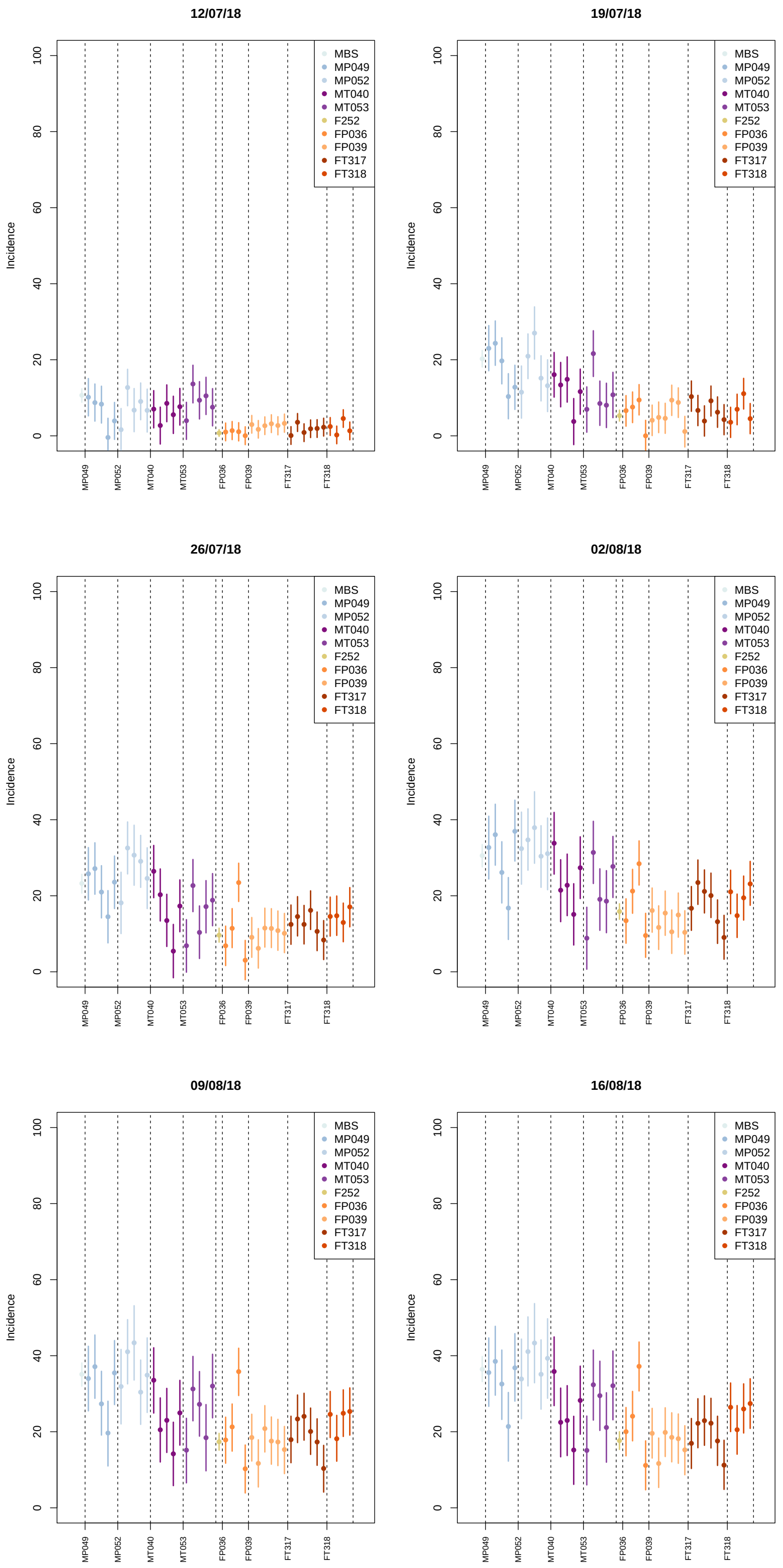

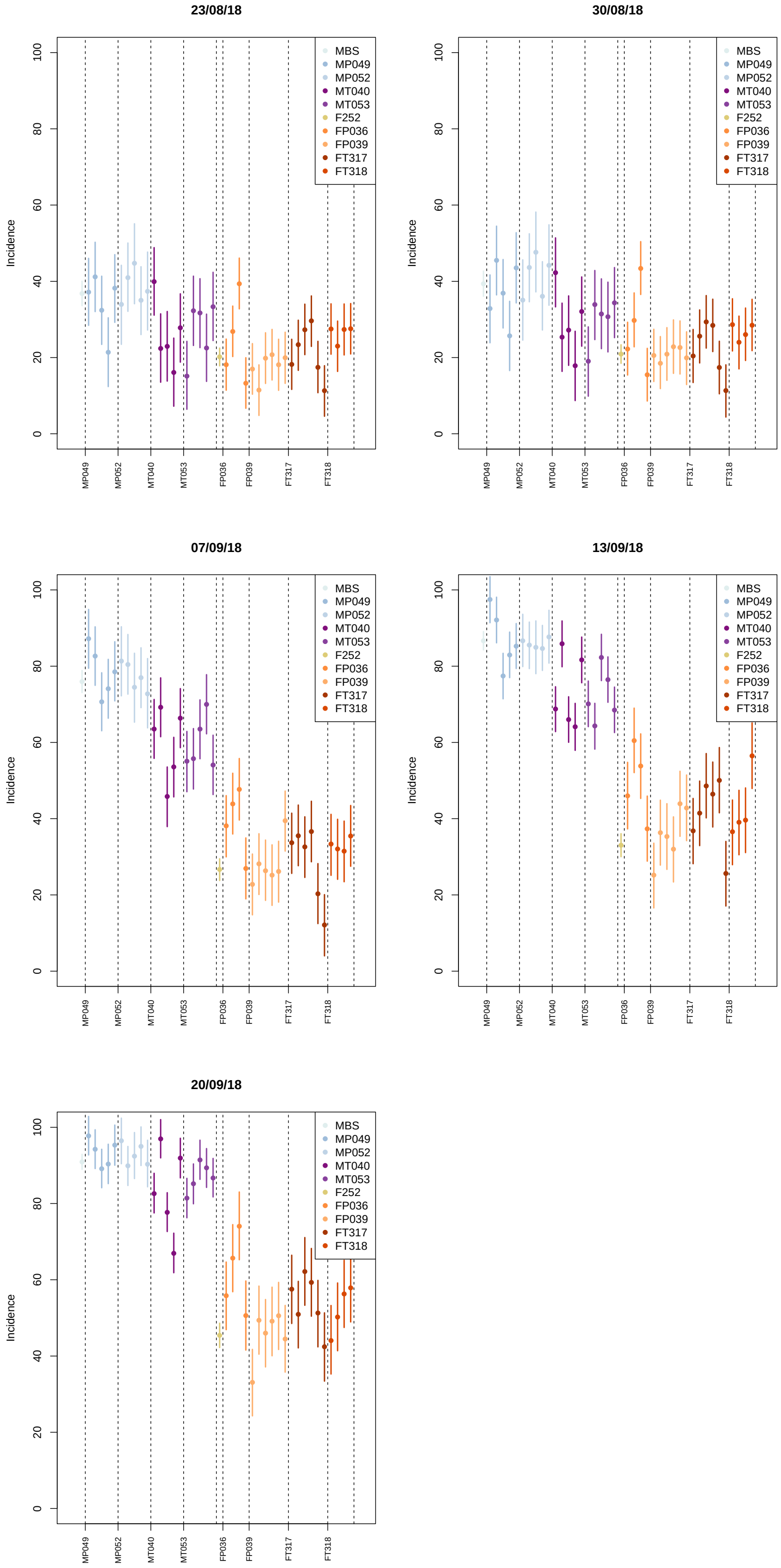

Figure S8 : Estimated medians of the ECB field incidence for each progenitor, the MBS847 control, and the F252 control at each monitoring date in 2018. Error bars represent 95% credible intervals for estimates.

### Supplementary Figure S9

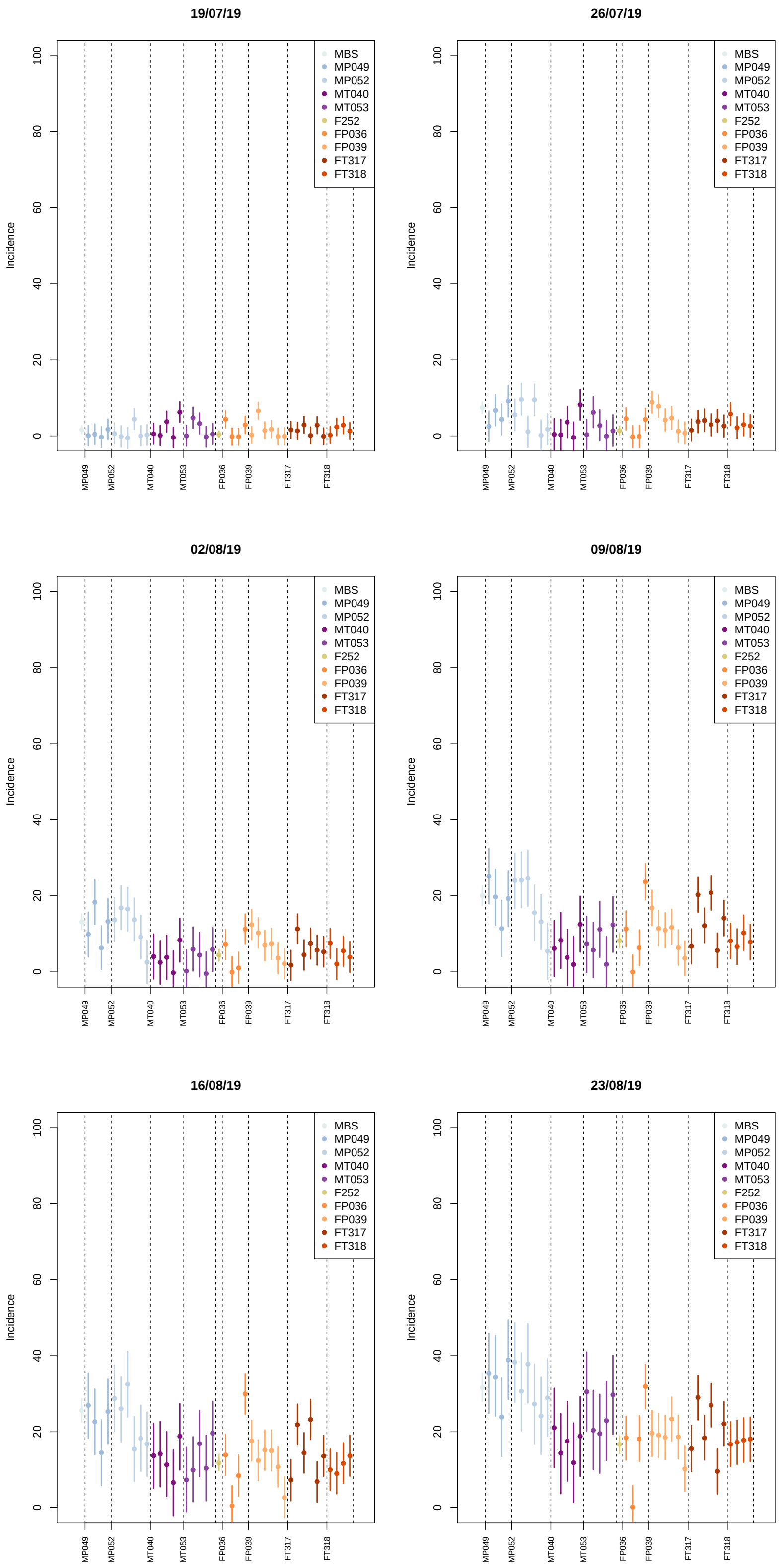

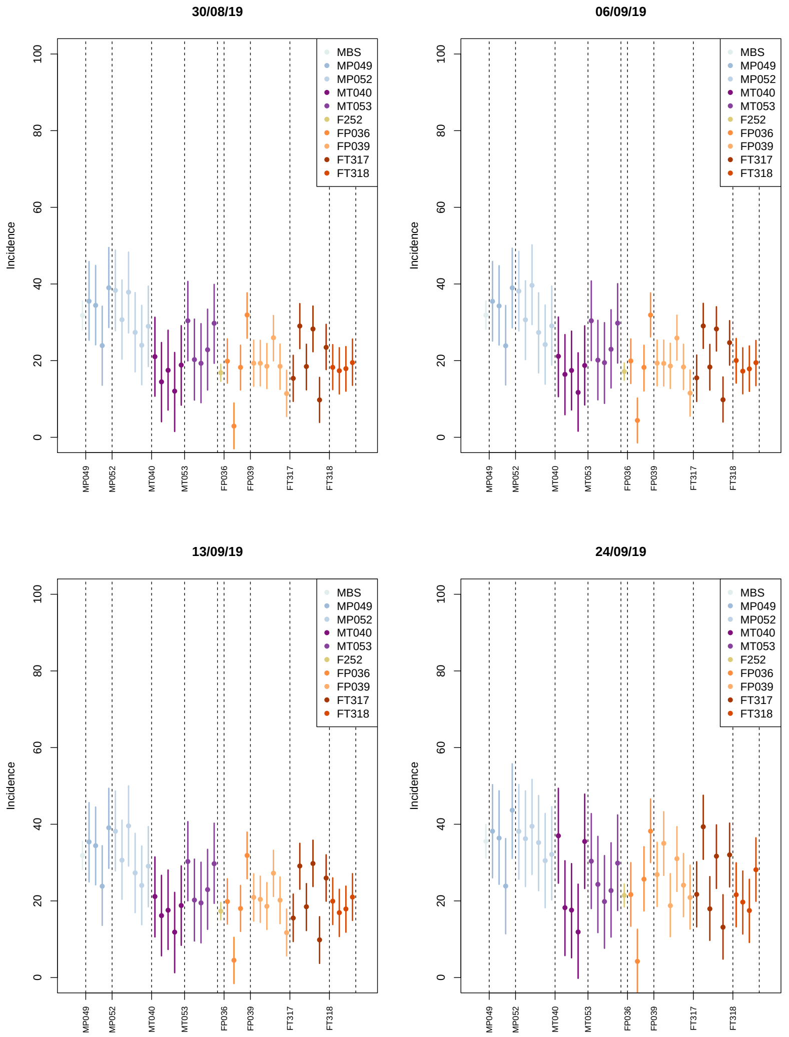

Figure S9 : Estimated medians of the ECB field incidence for each progenitor, the MBS847 control, and the F252 control at each monitoring date in 2019. Error bars represent 95% credible intervals for estimates.

### Supplementary Table 1

| Components | Abreviation | Unit |
|---|---|---|
| Median flowering | flo | thermal days (number of days at 20°C) |
| Median plant height | mhau | cm (from the soil to the base of the tassel) |
| Total weight | pdstot | g (fresh plant weight) |
| Weight of green parts | pdsgreen | g (fresh plant weight) |
| Mean ears weight | mpdsel | g |
| Dry matter | ms | percentage (100 x (dry weight/fresh weight)) |
| Cell Wall Residue | cwr | %ms (percentage of dry matter) |
| Silica content | cendre_ms | %ms (percentage of dry matter) |
| Klason lignin per cell wall residue | kl_cwr | %cwr (percentage of cell wall residue) |
| Klason lignin per dry matter | kl_ms | %ms (percentage of dry matter) |
| Acetyl bromid lignin | abl_cwr | %cwr (percentage of cell wall residue) |
| Acid Detergent Lignin | ADL | %ms (percentage of dry matter) |
| p-hydroxyphenyl sub-units | H | $\mu$moles/g Klason lignin |
| Guaiacyl sub-units | G | $\mu$moles/g Klason lignin |
| Siryngylic sub-units | S | $\mu$moles/g Klason lignin |
| Sum of lignin compounds | HGS | $\mu$moles/g Klason lignin (H+G+S) |
| Percentage p-hydroxyphenyl sub-units | pH | %HGS |
| Percentage of guaiacyl sub-units | pG | %HGS |
| Percentage of siryngylic sub-units | pS | %HGS |
| Syringyl/guaiacyl ratio | SoverG | number (S/G) |
| Neutral Detergent Fiber | NDF | %ms (percentage of dry matter) |
| KLADL_NDF | KLADL_NDF | %NDF (percentage of neutral detergent fiber) |
| Acid Detergent Fiber | ADF | %ms (percentage of dry matter) |
| Cellulose content per dry matter | cellulose_ms | %ms (percentage of dry matter) |
| Cellulose neutral detergent fiber | cellulose_NDF | %NDF (percentage of neutral detergent fiber) |
| Hemicellulose dry matter | hemicellulose_ms | %ms (percentage of dry matter) |
| Hemicellulose neutral detergent fiber | hemicellulose_NDF | %NDF (percentage of neutral detergent fiber) |
| PCAest | PCAest | mg/g CWR (per gram of cell wall residues) |
| Esterfied and Etherified Ferulic Acid | Fatot | mg/g CWR (per gram of cell wall residues) |
| Etherified Ferulic Acid | Faest | mg/g CWR (per gram of cell wall residues) |
| Estherified Ferulic Acid | Faeth | mg/g CWR (per gram of cell wall residues) |
| In vitro dry matter digestibility | IVDMD | %ms (percentage of dry matter) |
| In vitro cell wall residues digestibility | IVCWRD | %cwr (percentage of cell wall residue) |
| IVCWRDm | IVCWRDm | |

Table S1 : Plant traits and Biochemicals compounds measured in 2019 in the DSEs. Flowering time and plant height are the plot median value. Weights and biochemical composition result from the pooling of three plants per plot.

### Supplementary Table 2

| Background | Population | Family | TTF G22 (2018) | TTF G23 (2019) |
|---|---|---|---|---|
| MBS847 | Ancestral | x | 76,4 [76,1 - 76,7] | 87.7 [86,9 - 88.5] |
| MBS847 | Early | ME049 | 73,3 [72,4 - 74,1] | 84,1 [83,3 - 85,0] |
| MBS847 | Early | ME052 | 70,3 [69,4 - 71,1] | 83,6 [82,9 - 84,4] |
| MBS847 | Late | ML040 | 92,8 [92,0 - 93,6] | 100,8 [100,2 - 101,4] |
| MBS847 | Late | ML053 | 93,9 [93,2 - 94,7] | 100,8 [100,2 - 101,5] |
| F252 | Ancestral | x | 64,2 [63,9 - 64,5] | 75,4 [86,9 - 88,5] |
| F252 | Early | FE036 | 61,5 [60,6 - 62,4] | 69,4 [68,5 - 70,4] |
| F252 | Early | FE039 | 60,8 [60,0 - 61,6] | 71,0 [70,2 - 71,7] |
| F252 | Late | FL317 | 69,8 [69,1 - 70,6] | 83,9 [83,3 - 84,5] |
| F252 | Late | FL318 | 69,6 [68,8 - 70,4] | 84,4 [83,6 - 85,1] |

Table S2: Thermal time to flowering of the plant families of the DSE.

### Supplementary Table 3

| trait | $\mu$ | $\sigma^2$ | pvalback | pvalpop | pvalfam | pvalproge |
|---|---|---|---|---|---|---|
| medflo | 89.8819635 | 2.65375292 | * | *** | *** | - |
| H | 14.8456663 | 0.73558692 | *** | ** | * | - |
| pS | 47.1920969 | 0.93591651 | - | *** | - | - |
| SoverG | 0.9582423 | 0.03602941 | - | *** | * | - |
| pG | 50.2780972 | 0.94709394 | - | *** | - | - |
| NDF | 53.5433298 | 1.62202410 | - | *** | *** | - |
| ADF | 27.1131162 | 1.01688355 | - | *** | *** | - |
| cellulose_ms | 24.6937726 | 0.90558803 | - | *** | *** | - |
| cwr | 62.0038622 | 1.85996570 | - | *** | ** | - |
| IVDMD | 49.8499066 | 1.73211126 | - | *** | * | - |
| kl_ms | 9.3158160 | 0.38813705 | - | *** | - | - |
| ADL | 2.5469997 | 0.14675765 | - | *** | - | - |
| pH | 1.6883767 | 0.33596220 | *** | - | - | - |
| S | 286.9573915 | 18.31476908 | - | - | * | *** |
| cendre_ms | 6.9575932 | 0.66358110 | *** | - | - | - |
| ms | 31.4256351 | 1.90105007 | - | *** | *** | - |
| HGS | 614.5728497 | 45.70561526 | - | - | - | *** |
| Faeth | 2.6848540 | 0.04812755 | - | * | ** | - |
| G | 302.0551177 | 26.64818989 | - | * | - | *** |
| hemicellulose_ms | 26.5719128 | 0.87198315 | - | *** | ** | - |
| KLADL_NDF | 4.4788545 | 0.19837340 | *** | ** | - | - |
| pdstot | 1170.8111835 | 202.61324673 | - | * | *** | - |
| pdsgreen | 513.3740311 | 82.71322375 | *** | - | - | - |
| PCAest | 10.7854651 | 0.58962824 | ** | *** | - | - |
| hemicellulose_NDF | 49.7695540 | 0.80493110 | ** | - | - | - |
| abl_cwr | 16.7589314 | 0.41381836 | - | *** | - | - |
| IVCWRDm | 38.7628695 | 1.25343405 | *** | - | - | - |
| Faest | 4.3299506 | 0.24089708 | - | - | - | - |
| kl_cwr | 15.0047200 | 0.32851650 | ** | * | - | - |
| cellulose_NDF | 45.9060813 | 0.66145031 | * | - | - | - |
| Fatot | 6.8127971 | 0.26434832 | - | - | * | - |
| IVCWRDc | 20.6110465 | 1.28117622 | - | - | - | - |

Table S3: ANOVA results for the biochemical analysis. The mean ($\mu$) and residual standard deviation ($\sigma^2$) are indicated for each trait, as are the p-values of the genetic background (pvalback), population (pvalpop), family (pvalfam) and progenitor (pvalproge) effects. "***" indicates a significance level of $< 0.001$, "**" indicates $< 0.01$, and "*" indicates $< 0.05$.

### Supplementary Table 4

| Ctype | Ntype | pvalyear | pvalfam | pvalge | pvaldev | sigma | $R^2$ |
|---|---|---|---|---|---|---|---|
| A | F | 0.0411 | 0.3125 | 0.4049 | 6.56e-02 | 4.1638 | 0.1471 |
| A | F-E | 0.0515 | 0.3190 | 0.4071 | 2.88e-02 | 3.9734 | 0.1735 |
| A | F-E-D | 0.0522 | 0.3560 | 0.3842 | 2.17e-02 | 3.6448 | 0.1822 |
| A-B | F | 0.9990 | 0.2094 | 0.3431 | 8.79e-08 | 0.6954 | 0.4870 |
| A-B | F-E | 0.3130 | 0.2535 | 0.3651 | 2.23e-09 | 0.7854 | 0.5542 |
| A-B | F-E-D | 0.0656 | 0.2826 | 0.2599 | 8.61e-10 | 0.7148 | 0.5779 |
| A-B-C | F | 0.0483 | 0.3970 | 0.4741 | 4.49e-04 | 0.6828 | 0.2976 |
| A-B-C | F-E | 0.0050 | 0.4200 | 0.3505 | 2.22e-06 | 0.7441 | 0.4510 |
| A-B-C | F-E-D | 0.0002 | 0.5047 | 0.2893 | 1.67e-06 | 0.6908 | 0.4849 |

Table 1: Table S4: ANOVA results for different CNratio. Data were analyzed using the ANOVA model (**??**). The pvalues for the effect of the year (pvalyear), the family (pvalfam), the interaction between the year and the family (pvalge) and the developmental stage covariate are given for each possible CNratio, as well as the residual variance (sigma) and the corresponding Rsquare of the model (R2).

### 1 Supplementary Maths

*Modeling ECB incidence variations over the season*
A Gaussian model $Y_{kt} \sim N(\mu_{g(k)t}, \sigma^2)$ was used to estimate the the probability of attack by ECB in the field in which $Y_{kt}$ is the incidence of the $k^{th}$ plots of the field at week $t$ (1):

$$\mu_{g(k)t} = \alpha_{g_{(k)}} + \beta_t + (\alpha\beta)_{g_{(k)}t} + \phi_{kt}, \tag{1}$$

where $\alpha_{g(k)}$ denotes the progenitor or family regression parameters, $\beta_t$ the time regression parameters and $(\alpha \times \beta)_{g(k)t}$ their interactions. $\phi_{kt}$ denotes the spatio-temporal random effects modeled in ([**?**]) by (2) .

$$\begin{array}{l} \phi_t | \phi_{t-1} \sim N(\rho_T \phi_{t-1}, \tau^2 Q(W, \rho_S)^{-1}) \\ \phi_1 \sim N(0, \tau^2 Q(W, \rho_S)^{-1}) \end{array}, \tag{2}$$

with $\phi_t = (\phi_{1t}, .., \phi_{Kt})$ is the vector of random effects for time period $t$. The temporal auto-correlation is modeled by $\rho_T \phi_{t-1}$ and the spatial auto-correlation by the variance $\tau^2 Q(W, \rho_S)^{-1}$, $Q$ is defined in ([**?**]), again, the three spatial structures described above were tested. A Uniform prior is set for $\rho_S$ and $\rho_T$ and Inverse Gamma is set for $\tau^2$. For the season analysis, three independent chains were run for 350 000 iterations with a burning period of 50 000 and a thinning of 30.
